# Mechanics and energetics of walking and running up and downhill: A joint-level perspective to guide design of lower-limb exoskeletons

**DOI:** 10.1101/2020.04.07.029579

**Authors:** Richard W. Nuckols, Kota Z. Takahashi, Dominic J. Farris, Sarai Mizrachi, Raziel Riemer, Gregory S. Sawicki

## Abstract

Lower-limb wearable robotic devices can provide effective assistance to both clinical and healthy populations; however, how assistance should be applied in different gait conditions and environments is still unclear. We suggest a biologically-inspired approach derived from knowledge of human locomotion mechanics and energetics to establish a ‘roadmap’ for wearable robot design. In this study, we characterize the changes in joint mechanics during both walking and running across a range of incline/decline grades and then provide an analysis that informs the development of lower-limb exoskeletons capable of operating across a range of mechanical demands. Eight subjects (6M,2F) completed five walking (1.25 m ^-1^) trials at −15%, −10%, 0%, 10%, and 15% grade and five running (2.25 m s^-1^) trials at −10%, −5%, 0%, 5%, and 10% grade on a treadmill. We calculated time-varying joint moment and power output for the ankle, knee, and hip. For each gait, we examined how individual limb-joints contributed to total limb positive, negative and net power across grades. For both walking and running, changes in grade caused a redistribution of joint mechanical power generation and absorption. From level to incline walking, the ankle’s contribution to limb positive power decreased from 44% on the level to 28% at 15% uphill grade (*p* < 0.0001) while the hip’s contribution increased from 27% to 52% (*p* < 0.0001). In running, regardless of the surface gradient, the ankle was consistently the dominant source of lower-limb positive mechanical power (47-55%). In the context of our results, we outline three distinct use-modes that could be emphasized in future lower-limb exoskeleton designs 1) Energy injection: adding positive work into the gait cycle, 2) Energy extraction: removing negative work from the gait cycle, and 3) Energy transfer: extracting energy in one gait phase and then injecting it in another phase (*i*.*e*., regenerative braking).

## Introduction

Lower-limb robotic exoskeletons can apply assistive torque to reduce the metabolic energy used by biological muscles to produce the force and work for locomotion [1]. A majority of these successful exoskeletons have focused on providing assistance at the ankle within a laboratory setting [2-10]. More recently, devices have begun to move outside of laboratory confinement. Fully-autonomous, portable devices have been demonstrated to reduce the metabolic cost of walking with [11] and without [3] additional load and during running [12]. A key factor for all of these systems is the coordination between the wearable robot and the human user.

Researchers have dedicated significant time and effort to understanding the interaction between exoskeleton control strategies and the physiological response of the human user. The high-level method for generating control commands [13, 14], the shape, the timing and magnitude of the torque assistance profile [15-17], and the lower-limb joint where assistance is targeted [17-20] can all influence how the well the user responds. Notably, to date most exoskeleton research studies have focused on optimizing controllers for a single gait at a fixed speed on level ground. However, although great advances have been made by this approach, to fully parameterize control strategies for the real-world, a more diverse range of locomotor scenarios must be explored. While brute force parameter sweeps and human-in-the loop optimization have been used to determine torque profiles on an individual basis [6, 7, 10, 21], discovering an optimal policy can take many hours and may not generalize beyond current test conditions. Thus, there is a need for simpler approaches to exoskeleton control that do not rely on (re)tuning, but rather use insights into the mechanism of these tasks in order to be effective across variable locomotion conditions (*e*.*g*., speed, grade, gait).

As exoskeletons become increasingly mobile, a clear problem arises: How can engineers deliver systems that can assist in natural environments where locomotion involves adjusting speed, changing gait from walk to run, and moving uphill or downhill? Indeed, few exoskeleton studies have focused on incline/decline walking [4, 22] or compared assistance strategies across speeds [10] in which mechanistic explanations for performance outcomes were provided. Injection of positive power has been shown to be a promising approach for achieving metabolic cost reduction [23]; however, whether this approach is effective across all leg joints or if it is effective across different grades or gaits is unknown. We suggest that a bio-inspired mechanistic understanding of how people move and exchange energy between their lower-limb joints and the external environment is crucial for successful designs that make exoskeletons truly effective in real-world conditions.

In fact, this mechanistic approach has been previously applied to exoskeleton development and logically explains why the field has so heavily focused on the ankle as a target for assistance in level walking [2, 5, 9]. The ankle provides the majority of power on level ground [24] and disrupted ankle mechanics common in clinical populations make it a good target for assistance [25, 26]. Guidance from baseline human gait data has motivated a bioinspired approach to borrow ‘best’ concepts from the biological system to guide design of wearable devices. For example, our previous work to design and test a clutch-spring ankle ‘exo-tendon’ [5, 9, 10] was directly inspired by insights from imaging research examining ankle muscle-tendon interaction dynamics [27, 28].

The same mechanistic approach can be applied towards the development of exoskeletons in non-level gait. In moving to inclines and declines, fundamental physics shape mechanical demands on the legs. Muscles must add or remove net mechanical energy lost or gained according to changes in height of the center of mass (COM) [29, 30] and numerous studies have contributed to our understanding of the dynamics of uphill and downhill gait at various speeds [31-41]. Joint level mechanical analyses through inverse dynamics have provided more detailed insight into the sources of mechanical energy generation/dissipation moving uphill/downhill, respectively. In general, hip moments increase during incline walking to add net mechanical work; and knee moments increase during decline walking to subtract net mechanical work [32, 37, 38]. In incline running, the required increase in energy also results from a shift in net power output to the hip [31, 38]. Inverse dynamics analysis has also been used to evaluate the effect of aging on the joint kinematics and kinetics of uphill walking and reveals that older adults perform more hip work and less ankle work in both level ground and incline walking [34]. Other studies have demonstrated that individual joint dynamics can be used as a predictive tool for estimating the metabolic cost of walking, with 89% of the added metabolic cost of incline walking explained through changes in joint kinematics and kinetics [33].

The purpose of this study was to characterize changes in lower-limb joint mechanics during both walking and running across a range of incline/decline grades and then provide an analysis that informs lower-limb exoskeleton development (Fig. 1). More specifically, we sought to add an applied twist to current basic science understanding by focusing interpretation of the measured changes in human joint mechanics to guide the development of versatile exoskeleton systems with the ability to inject (net positive work), remove (net negative work) and transfer (net zero work) mechanical energy to meet variable mechanical demands of real-world environments.

**Figure 1:**
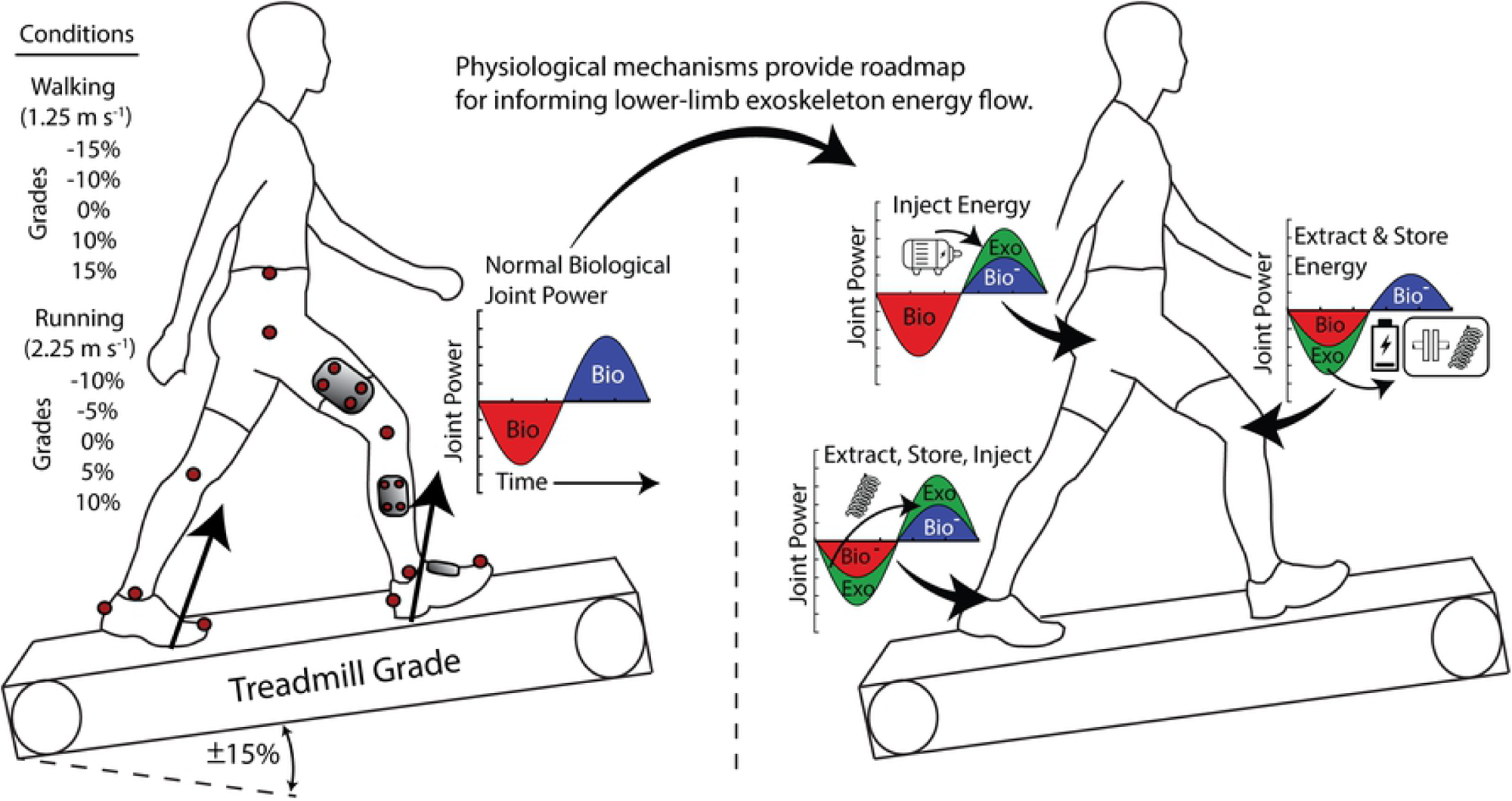
Schematic of experimental design and analysis. Representation of gait conditions for characterizing changes in lower-limb mechanics during walking and running across incline and decline grades. Example of energy cycle and potential mechanisms for how physiological mechanisms may provide a roadmap for informing lower-limb exoskeleton development.

## Methods

Eight adults (6M,2F, age: 23.38±4.10 yrs; mass 75.39±11.57 kg; height 177±0.07 cm) participated in the study. All subjects were healthy and gave written informed consent to participate in the study. The protocol and all testing were approved by the University of North Carolina at Chapel Hill Institutional Review Board.

Subjects completed five walking (1.25 m/s) and five running (2.25 m/s) trials over a range of incline and decline grades (Fig. 1). Walking trials were at −15%, −10%, 0%, 10%, and 15% and running trials were at −10%, −5%, 0%, 5%, and 10%. The ranges provided an overlap at the −10%, 0%, and 10% grade for comparison between the two gaits. All experimental trials took place on a split belt instrumented treadmill capable of incline and negative velocity (Bertec, Columbus, OH, USA). Decline gait was obtained by inclining the treadmill and reversing the belt velocity. Walking and running trials each lasted 7 minutes to ensure steady-state metabolic data. Walking and running trials were pseudorandomized, and once the treadmill incline was set, all conditions for that grade were completed.

Joint kinematic data were recorded using an eight camera motions capture system (VICON, Oxford, UK) to record the position of 22 reflective markers on the right lower limb and pelvis. Raw marker positions were filtered using a 2^nd^ order, low pass filter with a cut off frequency of 10 Hz. Segment tracking was performed by placing rigid plates containing clusters of 3-4 markers on the foot, shank, thigh, and pelvis. Calibration landmarks and relative location of tracking markers were identified through a standing trial that was performed at the beginning of the trials. The tracking markers were recorded during each trial and the orientation of the distal segment relative to the proximal segment was used to define the 3D joint angle. Ground reaction force (GRF) data was captured through the force plates embedded in the instrumented treadmill (BERTEC, Columbus, OH, USA). GRF data were filtered with a 2^nd^ order low pass Butterworth filters with a cut off frequency of 35 Hz.

The GRF and the kinematic data from the individual limbs were used to perform an inverse dynamics analysis. We performed inverse dynamics at the joint level using commercially available software (Visual 3D, C-motion, USA). Calculations of the time-varying moment and power were performed at the ankle, knee, and hip. Average positive and negative power (W kg^-1^) was calculated for each joint at each condition. Average positive power for each joint over the stride was calculated by integrating periods of only positive joint power with respect to time. This positive joint work (J kg^-1^) was then averaged across all of the strides. Average joint positive mechanical power was calculated by dividing the average joint positive work by the average stride time for the trial. The total limb average positive power was calculated by summing the average positive power at each joint total = hip + knee + ankle). Next, each individual joint’s percent contribution to the total limb average positive power for the stride was calculated by dividing that joint’s average positive power by the total limb average positive power. The same process was followed to compute stride average negative power, where only the contribution of negative work at each joint was used. The average net power for each joint and for the limb was then calculated by summing the positive and negative average power values at each joint and for the limb.

Whole body metabolic energy expenditure was captured using a portable metabolic system (OXYCON MOBILE, VIASYS Healthcare, Yorba Linda, CA, USA). Rates of oxygen consumption and carbon dioxide production during trials were recorded and converted to metabolic powers using standard equations [42]. Baseline quiet standing metabolic rate was captured prior to gait trials. For each condition, respiratory data from minute 4 to 6 were averaged and used to report the steady state metabolic energy consumptions (watts) for the trial. The metabolic system reported values that were averaged over 30 second intervals so four values were averaged for each trial. In the most extreme case of 10% incline running, subjects could not complete the trial while maintaining a respiratory exchange ratio (RER) below one. Therefore, only data from 3 out of 7 subjects are included for the 10% incline running condition. Task dependent metabolic power was calculated by subtracting the metabolic power in standing from the metabolic power recorded during the trial. These values were then normalized to each individual’s body mass. We then calculated cost of transport (COT) (J m^-1^ kg^-1^) by dividing mass normalized net metabolic power (W kg^-1^) by walking speed (m s^-1^):

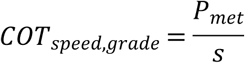

where *P*_*met*_ is mass normalized net metabolic power, and s is speed. Additionally, efficiency was calculated as the ratio of average total limb positive mechanical power to net metabolic power:

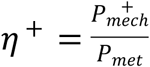

where *η*^+^ is efficiency of positive work, 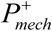 is the average total limb positive power (summed across the lower-limb joints), and *P*_*met*_ is mass normalized net metabolic power.

For each gait (walk and run), we performed a repeated measures ANOVA (rANOVA, main effect: grade) to test the effect of grade on stride average joint power of the ankle, knee, and hip. (α = 0.05; JMP Pro, SAS, Cary, NC). In addition, for each gait (walk and run), we performed a repeated measures ANOVA (rANOVA main effect: joint) to evaluate the relative contribution of each joint at each grade. We applied a post-hoc Tukey HSD (HSD) test to evaluate for significance between conditions (either grade or joint). Finally, we performed matched pair t-test to evaluate the effect of gait (walk, run) on the stride average joint power contributions for similar grades (−10%, 0%, and 10%). We did not run statistical analysis on metabolic data.

## Results

### Mechanical Power in Walking

#### Net Power

The average net mechanical power delivered at the ankle, knee, and hip all increased with grade (Fig. 2A). The average net power of the ankle increased with grade (rANOVA, *p* < 0.0001), was negative for decline conditions, and positive for level ground and incline grades. The average net power of the knee was negative in all conditions except the +15% grade. The knee was the largest source of net negative power in all conditions, and the magnitude increased as grade decreased (rANOVA, *p* < 0.0001). The average net power of the hip was positive in all conditions and increased with grade (rANOVA, *p* < 0.0001). As incline increased, we observed an increased reliance on the hip for the required net positive power.

**Figure 2:**
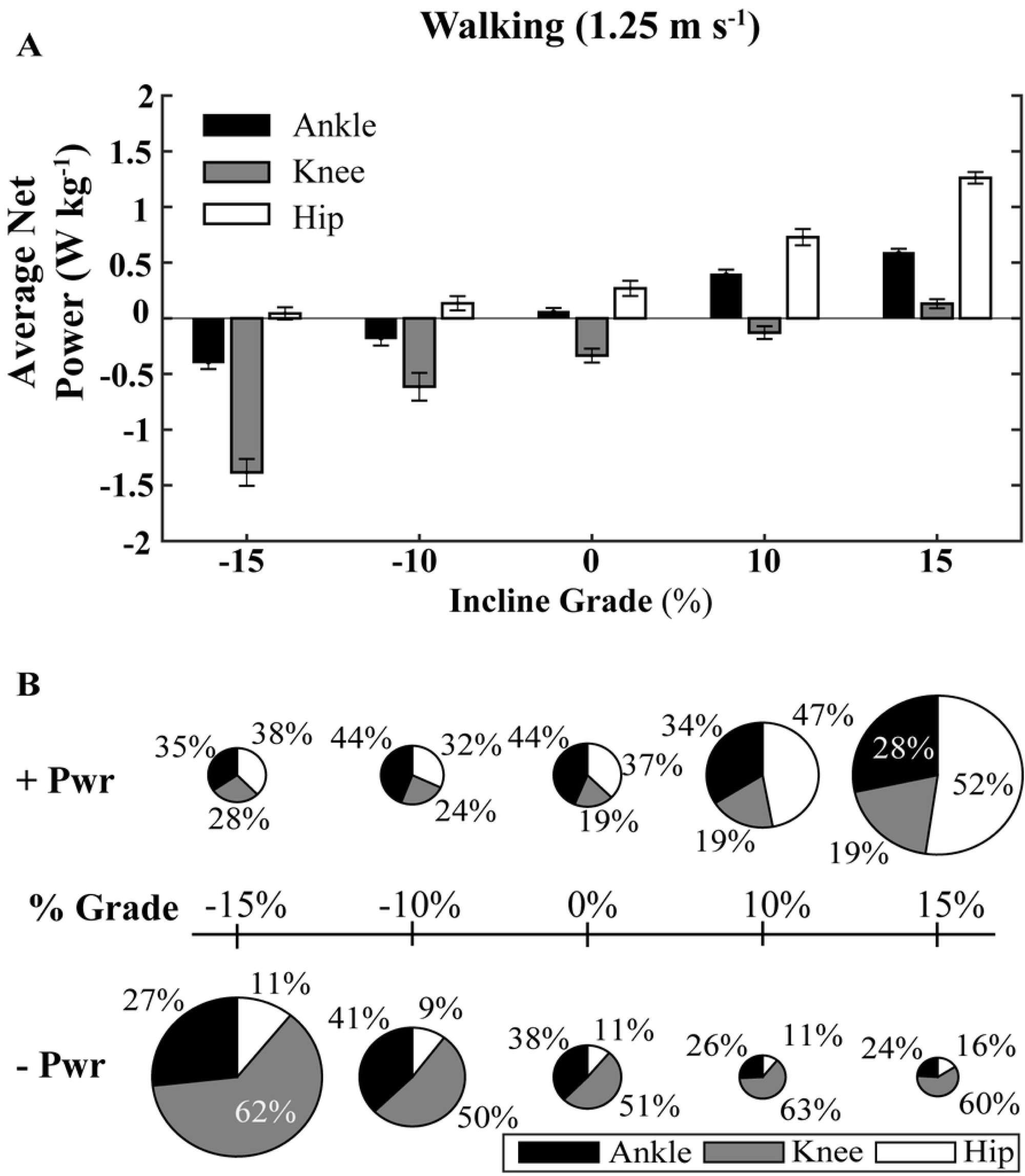
Percent distribution of average positive and negative lower-limb joint power for walking at 1.25 m s^-1^ over a range of grades. **(A) A**verage net power of each joint across surface grade conditions for walking. **(B)** The area of each pie is normalized to the average positive power at level grade for walking (1.02 W kg^-1^).

#### Positive Power

The average positive power of the limb (ankle + knee + hip) increased with increasing grade (rANOVA, *p* < 0.0001) (Table 1; Fig. 2B) from 1.02 W kg^-1^ at level to 1.70 W kg^-1^ (HSD, *p* < 0.0001) and 2.60 W kg^-1^ (HSD, *p* < 0.0001) at 10% and 15% grades respectively. Limb positive power was not significantly different from level at −10% and −15% grades respectively. The positive power of all three joints also increased individually with increased grade (rANOVA, *p* < 0.0001) (Table 1). However, the relative contribution of the ankle, knee, and hip to the total positive power of the limb changed with grade due to the unequal modulation of positive power at each joint for each grade (Table 2; Fig. 2B). In level walking, the ankle was the largest contributor to positive mechanical power at 44%, followed by 37% from the hip, and 19% from the knee (rANOVA, *p* = 0.0001; HSD, *p* < 0.0001). As grade increased, the percent contribution of the ankle decreased (rANOVA, *p* < 0.0001) to 34% at 10% grade (HSD, *p* = 0.0095) and 28% at 15% grade (HSD, *p* < 0.0001) relative to level. Conversely, the percent contribution of the hip increased with grade (rANOVA, *p* < 0.0001) from 37% at level to 47% at 10% grade (HSD, *p* = 0.0233) and 52% at 15% grade (HSD, *p* < 0.0001). For incline grades, the relative contribution of the knee to positive power was the smallest (19%) and did not change as the power was redistributed primarily between ankle and hip. For decline grades, the only significant shift in percent contribution to positive power was a decrease in the ankle contribution from 44% at level to 34% at −15% grade (rANOVA, *p* < 0.0001; HSD, *p* = 0.0167). There was no significant difference in the contribution to positive power among the joints at - 15% grade.

**Table 1:** Lower-limb joint average mechanical power for walking and running at multiple grades.

|  | Grade (%) | Joint Positive Power (W kg<sup>-1</sup>) |  |  |  | Joint Negative Power (W kg<sup>-1</sup>) |  |  |  |
| --- | --- | --- | --- | --- | --- | --- | --- | --- | --- |
|  |  | <u>Ankle</u> | <u>Knee</u> | <u>Hip</u> | <u>Total</u> | <u>Ankle</u> | <u>Knee</u> | <u>Hip</u> | <u>Total</u> |
| **Walk**<br>(1.25 m s<sup>-1</sup>) | -15 | 0.30 | 0.24 | 0.32 | 0.86 | -0.70 | -1.62 | -0.28 | -2.60 |
|  | -10 | 0.41 | 0.22 | 0.30 | 0.94 | -0.60 | -0.84 | -0.16 | -1.60 |
|  | 0 | 0.45 | 0.19 | 0.38 | 1.02 | -0.39 | -0.53 | -0.11 | -1.03 |
|  | 10 | 0.58 | 0.32 | 0.81 | 1.71 | -0.18 | -0.45 | -0.08 | -0.71 |
|  | 15 | 0.74 | 0.50 | 1.36 | 2.60 | -0.15 | -0.37 | -0.10 | -0.62 |
| **Run**<br>(2.25 m s<sup>-1</sup>) | -10 | 1.28 | 0.61 | 0.75 | 2.64 | -1.12 | -2.40 | -0.37 | -3.88 |
|  | -5 | 1.54 | 0.69 | 0.91 | 3.14 | -0.98 | -1.98 | -0.29 | -3.25 |
|  | 0 | 2.01 | 0.64 | 1.02 | 3.66 | -1.13 | -1.83 | -0.15 | -3.12 |
|  | 5 | 2.05 | 0.66 | 1.39 | 4.09 | -1.14 | -1.57 | -0.16 | -2.86 |
|  | 10 | 2.11 | 0.79 | 1.63 | 4.53 | -1.07 | -1.52 | -0.21 | -2.81 |

**Table 2:** Percent contribution of each joint to total limb power in walking at 1.25 m s^-1^. A repeated measures ANOVA (main effect: grade^##^) tested the effect of grade on stride average joint power of the ankle, knee, and hip (^#^ indicates HSD post-hoc comparison to 0% grade). In addition, a repeated measures ANOVA (main effect: joint*) evaluated the relative contribution of each joint at each grade. (main effect: joint \**p* = 0.0043; \*\**p* < 0.0001). Pairwise HSD was used to evaluate significant differences between joints.

| Joint Positive Power (W kg<sup>-1</sup>) |  |  |  |  |  |  |
| --- | --- | --- | --- | --- | --- | --- |
| Grade (%) | <u>Ankle</u> | <u>Knee</u> | <u>Hip</u> | <u>Pairwise HSD</u> |  |  |
|  | <sup>##</sup>*p* < 0.0001 | <sup>##</sup>*p* = 0.0203 | <sup>##</sup>*p* < 0.0001 | *Ank:Knee* | *Ank:Hip* | *Hip:Knee* |
| -15 | 34% | 28% | 38% |  |  |  |
|  | <sup>#</sup>*p* = 0.0167 |  |  |  |  |  |
| -10* | 43% | 25% | 32% | *p* = 0.0031 |  |  |
| 0** | 44% | 19% | 37% | *p* < 0.0001 |  | *p* = 0.0003 |
| 10** | 34% | 19% | 47% | *p* = 0.0001 | *p* = 0.0009 | *p* < 0.0001 |
|  | <sup>#</sup>*p* = 0.0095 |  | <sup>#</sup>*p* = 0.0233 |  |  |  |
| 15** | 29% | 19% | 52% | *p* = 0.0001 | *p* < 0.0001 | *p* < 0.0001 |
|  | <sup>#</sup>*p* < 0.0001 |  | <sup>#</sup>*p* < 0.0001 |  |  |  |
| Joint Negative Power (W kg<sup>-1</sup>) |  |  |  |  |  |  |
| Grade (%) | <u>Ankle</u> | <u>Knee</u> | <u>Hip</u> | <u>Pairwise HSD</u> |  |  |
|  | <sup>##</sup>*p* = 0.0077 | <sup>##</sup> - | <sup>##</sup>*p* = 0.0038 | *Ank:Knee* | *Ank:Hip* | *Hip:Knee* |
| -15** | 28% | 11% | 62% | *p* < 0.0001 | *p* = 0.0009 | *p* < 0.0001 |
| -10** | 41% | 9% | 50% |  | *p* = 0.0004 | *p* < 0.0001 |
| 0** | 38% | 11% | 51% | *p* = 0.0115 | *p* < 0.0001 | *p* < 0.0001 |
| 10** | 27% | 11% | 62% | *p* < 0.0001 | *p* < 0.0001 | *p* < 0.0001 |
|  |  |  | <sup>#</sup>*p* = 0.0433 |  |  |  |
| 15** | 24% | 16% | 60% | *p* < 0.0001 |  | *p* < 0.0001 |

#### Negative Power

The magnitude of stride average limb negative power decreased with increasing grade (rANOVA, *p* < 0.0001) from −1.03 W kg^-1^ in level to −0.73 W kg^-1^ at 10% grade (HSD, *p* = 0.1918) and −0.62 W kg^-1^ at 15% grade (HSD, *p* = 0.0305) (Table 1; Fig. 2B) Negative limb power was significantly larger in magnitude at −1.60 W kg^-1^ at −10% grade (HSD, *p* = 0.0015) and −2.60 W kg^-1^ at −15% grade (HSD, *p* < 0.0001). The knee contributed >50% to limb negative power, and the percent contribution was greater than that of the hip in all conditions and that of the ankle in all conditions but the −10% grade (rANOVA, p < 0.0001; HSD, *p* < 0.05) (Table 2; Fig. 2B). The percent contribution of the knee to negative limb power increased with incline (rANOVA, p = 0.0038) from 51% at level to 63% at 10% grade (HSD, *p* = 0.0433) and 60% at 15% grade and coincided with a decrease in ankle contribution (rANOVA, p = 0.0007). Ankle negative power contribution was maximized for −10% grade at 41%. Hip contribution to negative power did not change with grade and was 12% on average.

### Mechanical Power in Running

#### Net Power

Similar to walking, the stride average net power of each joint increased from negative to positive grade (rANOVA, *p* < 0.0001) (Fig. 3A). The average net power of the ankle and hip was positive in all conditions and increased in magnitude with increasing grade (rANOVA, *p* < 0.0001). In contrast, the average net power of the knee was negative in all conditions and became more negative in large downhill grades (rANOVA, *p* < 0.0001).

**Figure 3:**
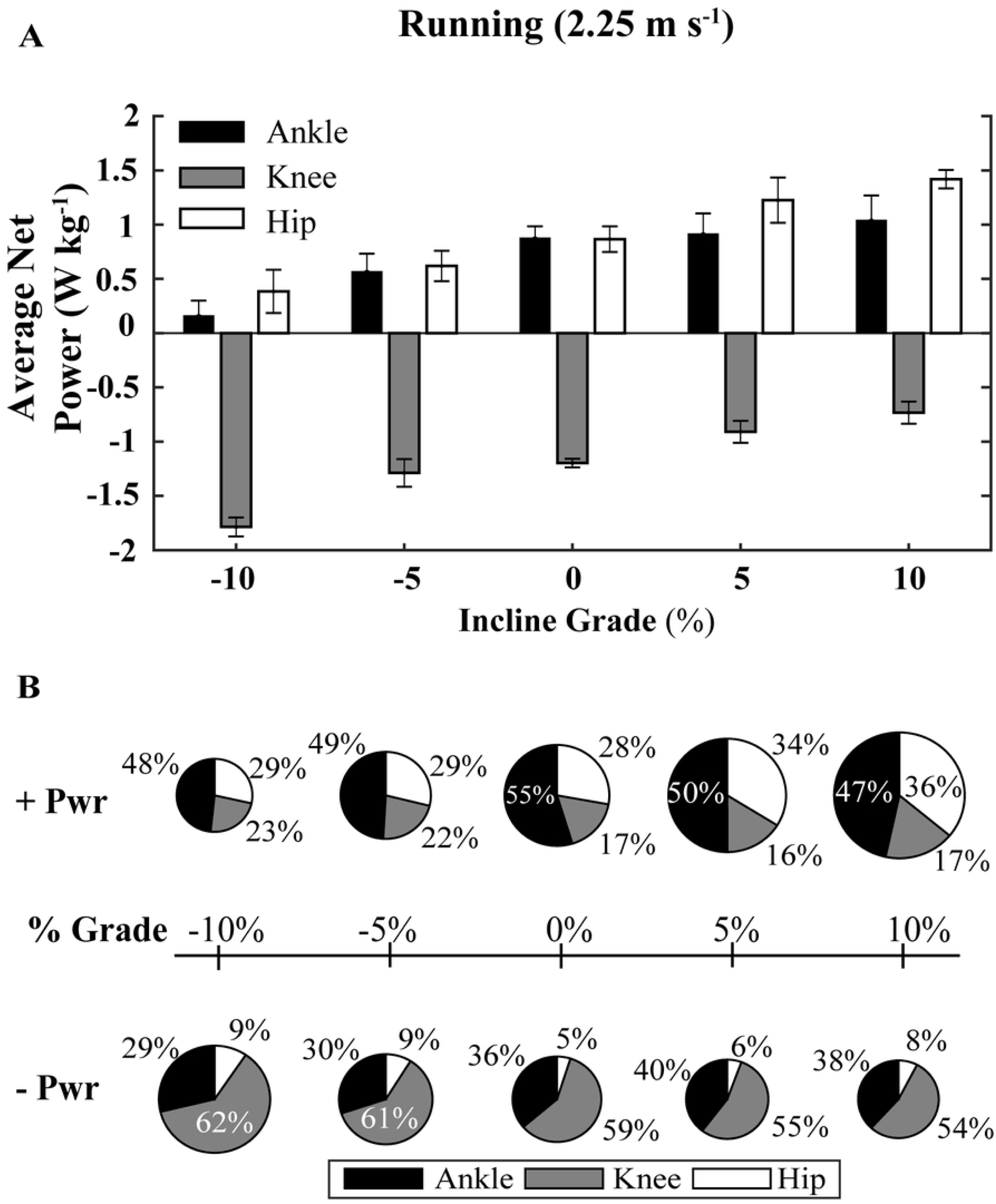
Percent distribution of average positive and negative lower-limb joint power for running at 2.25 m s^-1^ over a range of grades. **(A)** Average net power of each joint across surface grade conditions for running. **(B)** The area of each pie is normalized to the average positive power at level grade for running (3.66 W kg^-1^).

#### Positive Power

The average positive power of the limb (ankle + knee + hip) increased with increasing grade (rANOVA, *p* < 0.0001) (Table 1; Fig. 3B) from 3.66 W kg^-1^ at level to 4.12 W kg^-1^ and 4.53 W kg^-1^ (HSD, *p* = 0.0005) at 5% and 10% grades respectively. Limb positive power decreased to 3.14 W kg^-1^ at −5%, and to 2.64 W kg^-1^ (HSD, *p* < 0.0001) at −10% grade. The ankle was the dominant source of positive mechanical power (>46%) in all conditions and was significantly different from the knee (rANOVA, *p* < 0.0001; HSD, *p* < 0.0001) in all conditions and for the hip in all but the 10% grade (rANOVA, *p* < 0.0001; HSD *p* < 0.0171) (Table 3; Fig. 3B). With increasing incline, ankle positive power percent contribution decreased (rANOVA, *p* = 0.04) from 55% at level to 46% at 10% grade (HSD *p* = 0.0263) while hip contribution increased (rANOVA, *p* = 0.0032) from 28% to 36% in the level versus 10% grade condition (HSD, *p* = 0.0051). For decline grades, there was no significant shift in the joint positive power distribution.

**Table 3:** Percent contribution of each joint to total limb power in running at 2.25 m s^-1^. A repeated measures ANOVA (main effect: grade^##^) tested the effect of grade on stride average joint power of the ankle, knee, and hip (^#^ indicates HSD post-hoc comparison to 0% grade). In addition, a repeated measures ANOVA (main effect: joint*) evaluated the relative contribution of each joint at each grade. (main effect: joint \*\**p* < 0.0001). Pairwise HSD was used to evaluate significant differences between joints.

| Joint Positive Power (W kg<sup>-1</sup>) |  |  |  |  |  |  |
| --- | --- | --- | --- | --- | --- | --- |
| Grade (%) | <u>Ankle</u> | <u>Hip</u> | <u>Knee</u> | <u>Pairwise HSD</u> |  |  |
| | <sup>##</sup>$p = 0.04$ | <sup>##</sup>$p = 0.0032$ | <sup>##</sup>$p = 0.1468$ | *Ank:Knee* | *Ank:Hip* | *Hip:Knee* |
| -10** | 48% | 28% | 23% | $p < 0.0001$ | $p = 0.0002$ | |
| -5** | 49% | 29% | 22% | $p < .0001$ | $p = .0023$ | |
| 0** | 55% | 28% | 17% | $p < .0001$ | $p < .0001$ | $p = .0197$ |
| 5** | 50% | 33% | 16% | $p < .0001$ | $p = .0171$ | $p = .0186$ |
| 10** | 46%<br><sup>#</sup>$p = 0.0263$ | 36%<br><sup>#</sup>$p = 0.0051$ | 18% | $p < .0001$ | $p = .0013$ | |

| Joint Negative Power (W kg<sup>-1</sup>) |  |  |  |  |  |  |
| --- | --- | --- | --- | --- | --- | --- |
| Grade (%) | <u>Ankle</u> | <u>Hip</u> | <u>Knee</u> | <u>Pairwise HSD</u> |  |  |
| | <sup>##</sup>$p = 0.0027$ | <sup>##</sup>$p = 0.1109$ | <sup>##</sup>$p = 0.0094$ | *Ank:Knee* | *Ank:Hip* | *Hip:Knee* |
| -10** | 28% | 10% | 62% | $p < .0001$ | $p = .0003$ | $p < .0001$ |
| -5** | 31% | 9% | 60% | $p < .0001$ | $p < .0001$ | $p < .0001$ |
| 0** | 36% | 5% | 59% | $p < .0001$ | $p < .0001$ | $p < .0001$ |
| 5** | 41% | 5% | 54% | $p = .0495$ | $p < .0001$ | $p < .0001$ |
| 10** | 38% | 8% | 54% | $p < .0001$ | $p < .0001$ | |

#### Negative Power

The magnitude of limb negative power in running decreased with grade (rANOVA, *p* < 0.0001) from −3.12 W kg^-1^ at level to −2.86 W kg^-1^ and −2.81 W kg^-1^at 5% and 10% grade (Table 1; Fig. 3B). The limb negative power magnitude increased to −3.25 W kg^-1^ for −5% and to −3.88W kg^-1^ for −10% grade (HSD, *p* = 0.0002). Similar to walking, each joint contributed different amounts to total limb average negative power (rANOVA *p* < 0.0001) (Table 3; Fig. 3B). The knee was the dominant source of negative power, producing >54% for all conditions and contributed significantly more than the ankle or hip (HSD *p* < 0.0001). The ankle contributed approximately 35% of the stride average negative power across all grades and the hip contribution was minimal (∼7%).

### Comparisons of Walking to Running

The average limb positive power was greater in running than walking. Switching from walking to running on level ground resulted in an increase in the ankle’s percent contribution from 44% to 55% (paired t-test *p* = 0.0024) and a decrease in the hip’s percent contribution from 37% to 28% (paired t-test *p* = 0.0196). The trend was similar at 10% grade, where switching from walking to running resulted in an increase in the ankle’s percent contribution from 34% to 46% (paired t-test *p* = 0.0024) and a decrease in the hip’s percent contribution from 47% to 36% (paired t-test *p* = 0.0196). The transition from walking to running at the 10% grade resulted in the hip being replaced by the ankle as the dominant contributor to positive power. For negative power at the 10% grade, switching from walking to running resulted in an increase in the ankle’s percent negative contribution from 27% to 38% (paired t-test *p* = 0.001) and a decrease in the knee’s percent contribution from 62% to 54% (paired t-test *p* = 0.0338).

### Temporal component of power redistribution

Time series plots show the redistribution of joint moment and power over the stride cycle for walking (Fig. 4) and running (Fig. 5). Again, the general trend was a shift in positive power generation to the hip with increasing incline, while the knee was the primary site of negative work (i.e., absorption). Changes in ankle positive power were predominantly seen at push-off (∼60% stride), with changes in the knee negative power and hip positive power coming in initial stance.

**Figure 4:**
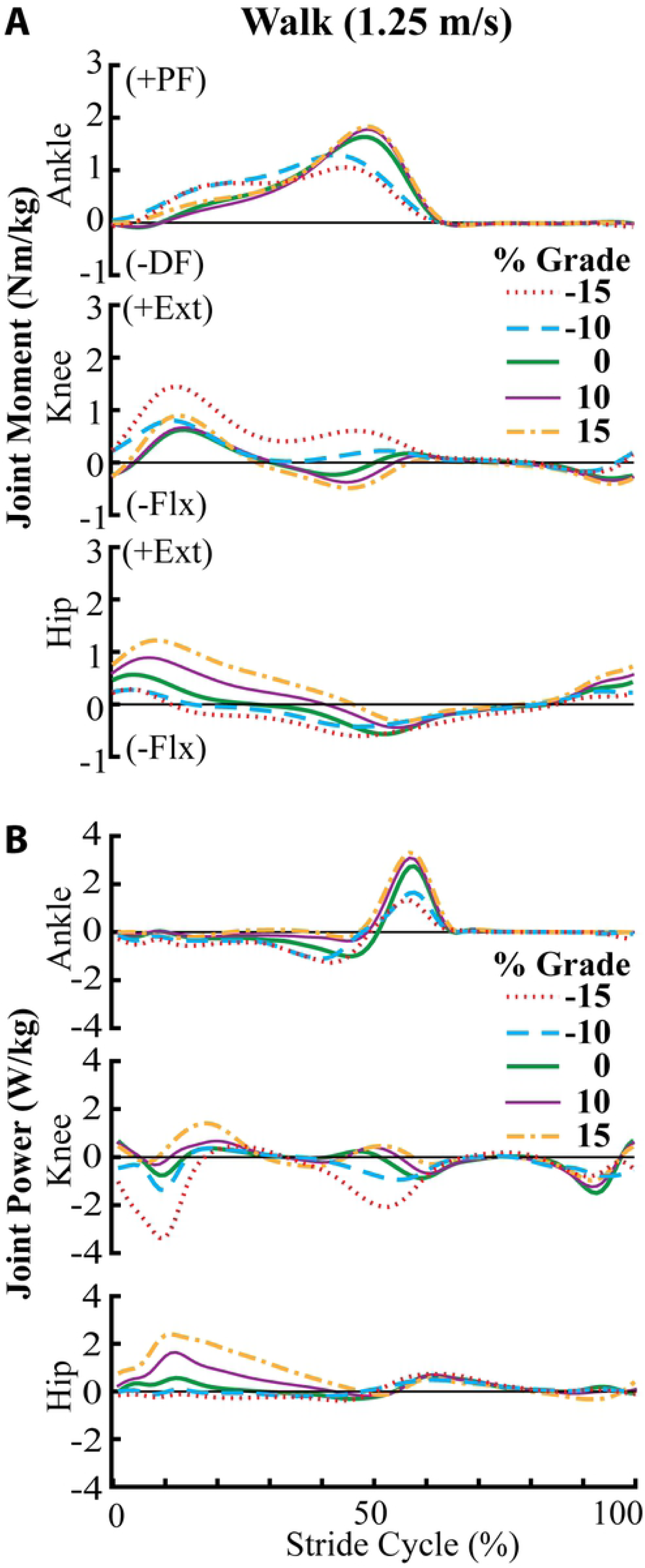
Ankle, knee, and hip joint kinetics for walking at 1.25 ms^-1^. Body-mass specific **(A)** joint moment (Nm kg^-1^) and **(B)** joint power (W kg^-1^) over a stride from heel strike (0%) to heel strike (100%) of the same leg for walking across surface grades from −15% downhill to +15% uphill.

**Figure 5:**
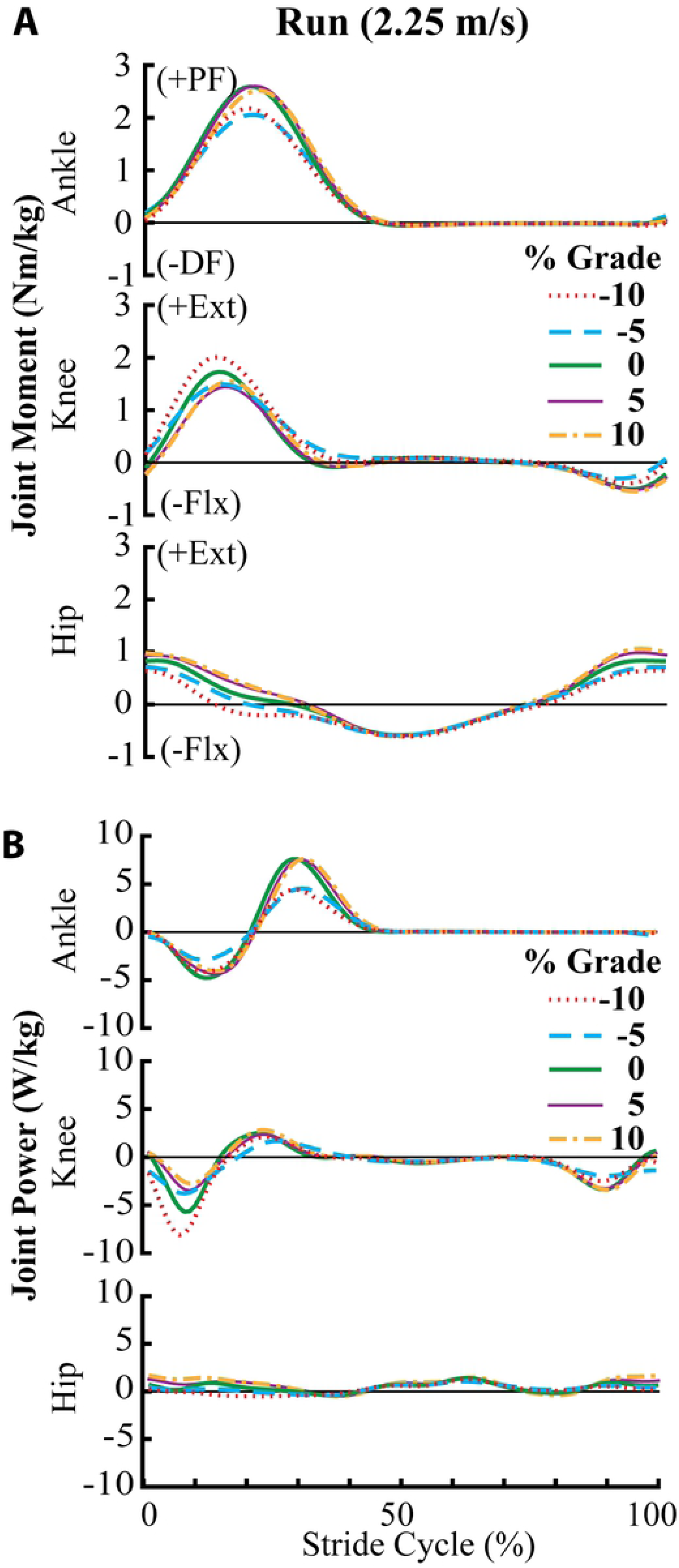
Ankle, knee and hip joint kinetics for running at 2.25 ms^-1^. Body mass-specific **(A)** joint moment (Nm kg^-1^) and **(B)** joint power (W kg^-1^) over a stride from heel strike (0%) to heel strike (100%) of the same leg for running across surface grades from at −10% downhill to +10% uphill.

### Metabolic Energy Demand

In walking, the measured metabolic minimum was at −10% grade (1.5 W kg^-1^) (Table 4). For running, the metabolic minimum was also at −10% grade (5.75 W kg^-1^) which was the steepest downhill grade tested in running. Efficiency of positive work was maximized at −10% grade in walking with an efficiency of 0.62.

**Table 4:** Net metabolic power, summed (ankle +knee +hip) lower-limb joint average positive power, efficiency of positive joint work, and cost of transport for walking and running up and downhill.

| | Grade (%) | $P_{\text{MET}}$ (W kg<sup>-1</sup>) | $P^+$ (W kg<sup>-1</sup>) | $\eta^+_{\text{WORK}}$ | COT (J kg<sup>-1</sup> m<sup>-1</sup>) |
| --- | --- | --- | --- | --- | --- |
| **Walk**<br>(1.25 m s<sup>-1</sup>) | -15 | 2.24 | 0.86 | 0.38 | 1.79 |
|  | -10 | 1.50 | 0.94 | 0.62 | 1.20 |
|  | 0 | 2.82 | 1.02 | 0.36 | 2.25 |
|  | 10 | 6.15 | 1.71 | 0.28 | 4.92 |
|  | 15 | 10.54 | 2.60 | 0.25 | 8.43 |
| **Run**<br>(2.25 m s<sup>-1</sup>) | -10 | 5.75 | 2.64 | 0.46 | 2.55 |
|  | -5 | 7.32 | 3.14 | 0.43 | 3.25 |
|  | 0 | 9.09 | 3.66 | 0.40 | 4.04 |
|  | 5 | 11.64 | 4.09 | 0.35 | 5.17 |
|  | 10 | 14.37 | 4.53 | 0.32 | 6.39 |

## Discussion

Our aim in this study was to measure and analyze human biomechanical response during walking and running on sloped surfaces in order to build a roadmap to help guide development of lower-limb wearable robots capable of adjusting to changing mechanical demands in real-world environments. We characterized the distribution of positive and negative mechanical power output across the lower-limb joints for incline and decline grades during walking and running. Our results confirm and are supported by previous studies demonstrating that the energetic demands of the lower limbs heavily depend on both ground slope and gait [31-41, 43-46]. Energy must be injected or extracted to raise or lower the potential energy of the center of mass (COM) for incline/decline walking.[29, 30]. Indeed, our data confirm that in both walking and running gait, the stride average total limb (ankle + knee + hip) power changes from net negative on decline grades to net positive on incline grades. Our findings also agree with previous work demonstrating the ankle to be a dominant source of positive mechanical power during both level walking and running gait [47], but that for incline walking the hip becomes an important source of positive mechanical power generation [31, 34, 35]. In addition, our data confirm that the knee is the dominant source of mechanical energy absorption during both walking and running across grades [38]. In the following sections we first discuss the biomechanical implications of our results and then focus on how these data could be utilized to create lower-limb wearable exoskeletons (or perhaps prostheses) that can respond to and perhaps even take advantage of changing mechanical demands across grades and gaits.

### Relationship between structure and function across task demand

The functional role of the ankle and the hip across grades aligns with the physiological structure of each joint’s muscle-tendon units (MTs). The hip MTs have short tendons and long muscle fascicles with low pennation [48]. In contrast, the structure of the ankle plantarflexor MTs, comprises relatively short, pennate muscle fibers in series with long compliant tendons. Added compliance in distal MTs make them ideal for storage and return of elastic energy during the gait cycle [48-50]. In incline gait, mechanical energy must be added to the body. Prior studies suggest that the structure of the MTs in the more proximal joints (*i*.*e*., hip) may be better suited to performing work on the COM because short, stiff tendons can directly transmit the work of the muscles to power the joint [48]. Furthermore, long muscle fascicles allow for production of force over a relatively larger range of motion and are important in incline walking due to larger joint range of motion.

In line with the idea that structure drives function, our walking data demonstrate a shift to power output in more proximal joints with an increase in incline. This finding is similar to prior studies which also show the dominant source of positive mechanical power shifts from the ankle to the hip in uphill walking [22, 32]. On the contrary, we found no evidence of a redistribution of positive work to the hip during uphill running. In running, the ankle still produced 46% of the positive power at 10% uphill grade. This finding seems to be in contrast with a previous study which showed that the hip contributed most to the increase in work for incline running [31]. However, our results may differ due to the different grade (6° and 12°), faster speed (3.0 and 3.5 m s^-1^), and lack of treadmill use in [31]. Interestingly, the ankle also performed a significantly higher percentage of the negative work in uphill running at 10% grade when compared to the level. This trend suggests that energy cycling through elastic mechanisms may still be an important feature retained in uphill running [51]. Due to the need for faster acceleration of the body in uphill running, ankle joint elasticity may facilitate higher peak powers and more net work output from the plantarflexors[48] by decreasing the required shortening velocity of the muscle fascicles of the ankle. Indeed, *in vivo* studies where ultrasound images of the triceps surae were taken in running and walking showed series elastic tissues allow the muscles to operate at lower average shortening velocities and that elastic recoil contributes substantially to positive work [28]. Additional *in vivo* studies of human muscle function, especially at proximal joints, in uphill and downhill walking and running would shed light on how MT architecture interacts with task demand for mechanical power generation /dissipation.

### Balance of positive and negative power varies across joint and grade

Net mechanical power production of the limb was governed by a balance between positive and negative power output that varied from joint to joint. The hip’s contribution to walking and running on sloped surfaces was net positive across all grades and gaits we tested and was modulated predominantly by changes in production of positive power (Tables 1-3, Figs. 2, 3). Despite large adjustments in net positive power output across grades, the hip was not the largest absolute contributor of positive power in most conditions (except incline walking). This was because the hip contributes very small amounts of negative power across conditions.

Conversely, the knee net power output was modulated predominantly by adjusting the production of negative power. In fact, the knee was the dominant contributor (>50%) to negative power across all grades in both walking and running. In all except the highest incline walking grade, the knee produced more negative than positive power, resulting in negative net power.

At the ankle, adjustments in lower-limb joint power production across grade/gait were more balanced in comparison to the hip (positive work modulated) and knee (negative work modulated). The average net power of the ankle was generated by adjustments to both positive and negative power across grade and gait. (Table 1, Figs. 2, 3) In level walking, the net power from the ankle was smaller than the hip despite the larger contribution to positive power from the ankle (Tables 1&2, Fig. 2). During incline walking, the ankle’s percent contribution to both positive and negative power decreased, potentially reflecting a reduced capacity to store and return elastic energy in the Achilles tendon. In decline walking, we observed the opposite trend where ankle net power was negative reflecting an increased capacity to store energy. In running, the ankle was the dominant source of positive mechanical power across all grades and the net power of the ankle was positive for all grades. (Table 1, 3, Fig. 3).

### Metabolic power and efficiency

Similar to Margaria *et al*. [30], we found that the greatest efficiency of positive work at - 10% slope for both walking and running. Additionally, the efficiency of positive work during walking at the extreme uphill (+15%) was ∼0.25 reflecting the efficiency of muscle-tendons during tasks exhibiting predominantly positive work [29, 52-55].

### Implications for lower-limb exoskeleton development

How the biological system distributes power across the joints in a variety of gait conditions has important implications for development of wearable assistive devices. To develop a roadmap for lower-limb exoskeleton design, we first define three main modes of operation: 1) (Net +) Energy injection – the device adds mechanical energy to the gait cycle using external sources of energy; 2) (Net -) Energy extraction – the device removes mechanical energy from the gait cycle to be dissipated as heat or stored (*e*.*g*., as mechanical energy in a spring or electrical energy in a battery); 3) (Net 0) Energy transfer – the device extracts energy at one time during gait and then injects it within or across joints at some time later (Fig. 6). With these modes the energy which is added, removed, or transferred may have different effects on the user’s biological and total joint power outputs, and, while most studies have a goal in mind (*e*.*g*., reduce biological moments and powers), the effects are often non-intuitive and hard to predict. Because the effect of an assistive device on the user is heavily dependent on the individual user’s biomechanical response, we further propose and discuss three potential biomechanical outcomes resulting from any of these modes of operation. The magnitude of the user’s biological joint power could: O1) decrease (=replacement) O2) remain constant (=augmentation), or O3) increase (=enhancement). Here, we offer several examples that span the possible physiological response outcomes (O1-3) for devices that inject positive power, but the same principles also apply for the other device modes as well (*i*.*e*., extraction and transfer).

**Figure 6:**
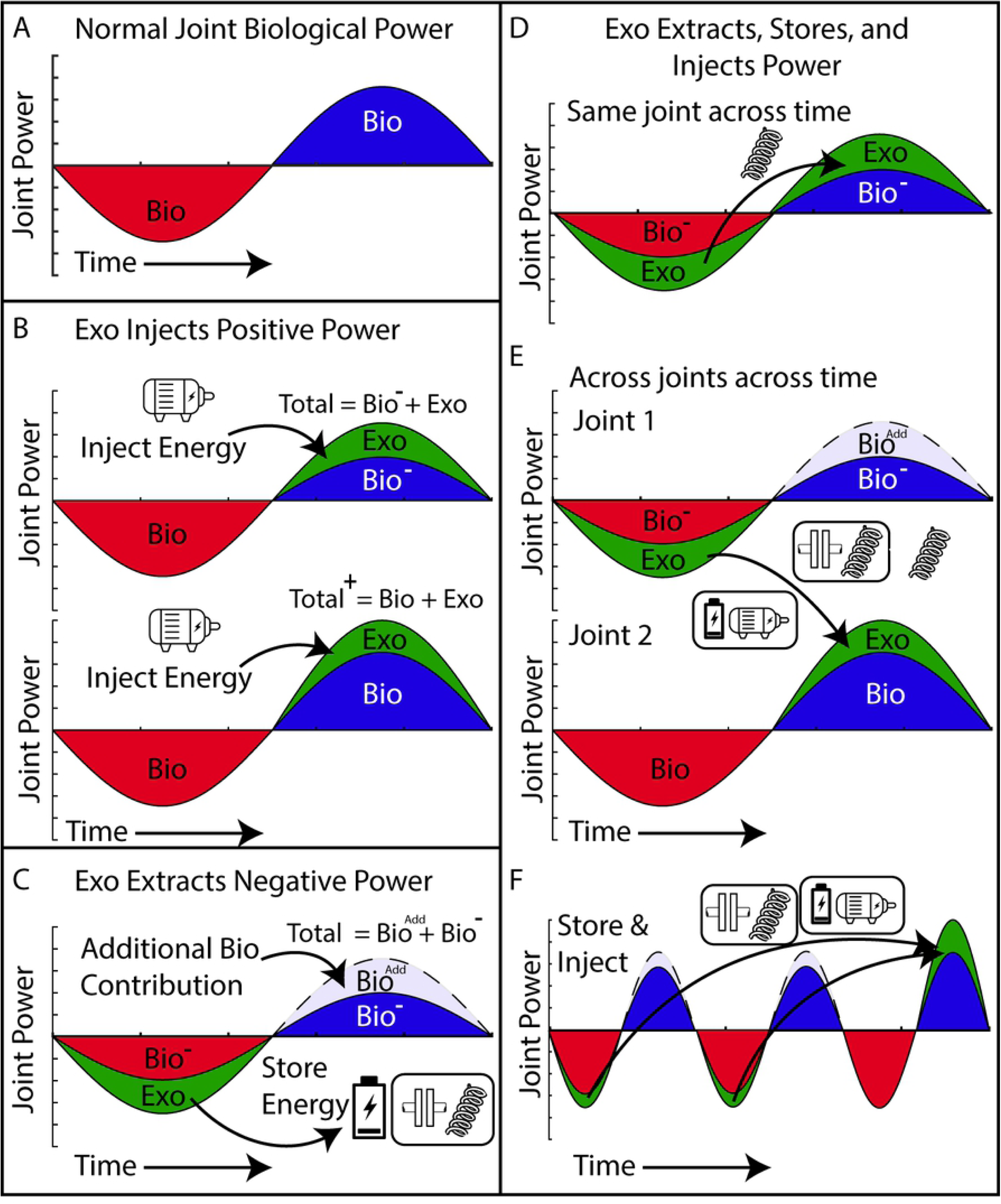
Potential mechanisms for exoskeleton energy exchange. **(A)** Example of energy cycle for a joint where negative joint power (red) is followed by positive joint power (blue) similar to the ankle power cycle during gait. **(B)** The exoskeleton (green) produces positive power and injects energy at the joint during the positive power phase of the gait via a motor or some other energy source. (Top) The positive bio power is reduced such that the total (bio+exo) positive power output of the joint remains the same (*i*.*e*., replacement). (Bottom) The additional energy increases the total (bio+exo) positive power output of the joint (*i*.*e*., augmentation). This is the most common mode employed on powered exoskeletons [3, 6, 7, 14, 17]. **(C)** The exoskeleton (green) produces negative power and extracts energy from the joint during the negative power phase of the gait via a damper or some other energy sink and, in this example, the user maintains the total (exo+ bio) negative power output of the joint, enabling a reduced biological contribution (*i*.*e*., replacement). In this mode, the exoskeleton negative power could drive an electrical generator and energy could be stored in a battery or used to power electronic devices [18, 56, 57]. If the negative power is normally recycled within the body and transferred to the positive power phase, additional biological power may be required to maintain biological positive power output (Bio^Add^). **(D-F)** The exoskeleton (green) could also operate in transfer mode by sequencing extraction and injection phases within or across the joints over time. **(D)** In the simplest case the exoskeleton stores energy during the negative power phase and returns it immediately to the same joint (*e*.*g*., with a spring) and, in this example, the user maintains the total joint power output enabling a reduction in both biological positive and negative power (=replacement) [5]. Other variants on transfer mode include: **(E)** The exoskeleton extracts energy at one joint (similar to C) and then immediately injects it at another (similar to B) [2]. **(F)** The exoskeleton extracts energy at one joint (*e*.*g*., with a spring or generator), temporarily stores it (*e*.*g*., using a battery or a clutch) and then after some delay injects it at the same joint (*e*.*g*., using a motor powered by the battery or spring recoil on release of a clutch).

#### Energy Injection

The first mode of device operation entails adding positive mechanical work at a joint(s) when the joint is producing positive power. This is the most prevalent strategy used in exoskeletons targeting the hip, knee, and ankle with the common desired goal being the reduction of metabolic demand in healthy individuals [3, 6-8, 14, 17, 23, 58]. The common expectation is the outcome where the addition of mechanical power causes a concomitant reduction of biological power while total power mostly remains constant (O1: replacement). While it’s been demonstrated that users will reduce biological moment such that the total joint moment remains invariant [59, 60], reductions in biological power often do not reflect full replacement [17, 61]. Thus, unlike what might be desired, the second physiological response outcome is often observed. Here, the biological power is reduced by less than the exoskeleton injects and the magnitude of the total joint power is increased (O2: augmentation) (Fig. 6B) [7]. [17, 61]. The third physiological response outcome is that the addition of exoskeleton positive power causes an enhancement of the biological power (O3: enhancement). It is possible that when injecting positive exoskeleton power, the user actually increases their biological power output and thus enhances the total joint power beyond the exoskeleton’s contribution. So far, we are not aware of cases where this physiological response has occurred, but it would be desirable for assistive and rehabilitative technology intended to improve function in clinical populations with baseline deficits in limb and joint power output (*e*.*g*., post-stroke) [62]. For example, the addition of positive power during push-off may help promote the recruitment of weak plantarflexors in stroke survivors or older adults. Studies have begun to demonstrate the potential for enhancing performance in clinical populations by providing positive power to the ankle [26, 63], however the actual effect on biological power is still unclear.

How might an engineer use employ the roadmap offered by this study to guide the strategy for exoskeleton positive power injection beyond level walking? The most notable example comes from the observed shift to hip dominated positive power in walking uphill (Figs. 2, 4). Given limited power supply of the device, our data would suggest that assistance should be redirected away from the ankle to the hip when transitioning to incline walking. Conversely, for running (Figs. 3,5), the ankle is the largest contributor to positive average power across *all* slopes and thus, shifting assistance to the hip may not be as beneficial.

#### Energy Extraction

The second mode of device operation involves removing negative mechanical work at a joint(s) when the joint is producing negative power. The extracted mechanical energy could be dissipated as heat (*e*.*g*., in a damper) or harvested to generate electricity which can then be stored in a battery or used to power electronic devices (Fig. 6C). Additionally, an exoskeleton that effective extracts energy from the gait cycle can potentially reduce the negative power required from muscles which, unlike many mechanical systems, require energy to elongate under load [64]. Similar to the effects from injecting positive power, generation negative power with exoskeletons may have a range of effects on the biological system that can be non-intuitive. For example, if an exoskeleton offloads a portion of the negative biological power at a joint, and that power was derived from stored energy in elastic tissues which can no longer be returned, it is possible that additional biological power may need to be generated in the positive phase to make up for lost energy stores (Fig. 6C). However, in the nominal case where the negative biological power is merely dissipated as heat rather than recycled, then the reduction in total power during the latter half of the cycle may not be problematic.

The knee has been the focus of energy harvesting exoskeletons due to its production of substantial negative power in gait, especially near the end of swing phase of walking (Fig. 4). There are several indications that if done correctly it is possible to generate electrical energy while reducing the muscle energetic demands and whole body metabolic cost [18, 56, 65, 66]. With consideration to changing mechanical demands on slopes surfaces, our results suggest enormous potential for harvesting energy using a knee exoskeleton during decline walking due to large increases in knee negative power throughout the gait cycle (Figs. 2, 4). In running, a knee exoskeleton may be widely versatile because the knee generates a large amount of negative power across all slopes including on inclines (Figs. 3, 5).

Although the ankle produces substantial negative power, harvesting exoskeletons might be ineffective in level gait because much of the joint power is recycled in elastic tissues [28], and thus as mentioned previously, the biological system would need to replace these losses with costly muscle work during a positive power phase at some joint in the limb. However, because ankle negative power increases and positive power decreases on declined surfaces (Fig. 2), energy harvesting may be a viable candidate at the ankle for decline walking.

#### Energy Transfer

The third mode of device operation is to transfer energy from one phase to another across the gait cycle either within or across joints (Fig. 6D-F). In this mode, because the exoskeleton extracts energy in the negative phase (*e*.*g*., Fig. 6C) and then injects the same energy later (*e*.*g*., Fig. 6B) in a positive phase, external power consumption of the device can be minimized (*e*.*g*., by using passive elements like springs and clutches) [67]. In addition, intra-joint transfer of energy from a negative power phase to a positive power phase may help mitigate the complication of the reduced biological power in the latter half of the power cycle. As depicted in Figure 5D, it is possible that the total power output of the joint (exo+bio) remains constant despite the reduction of biological power in both the negative and positive power phases. The simplest device applying this mode of operation is an elastic exoskeleton that uses a spring in a parallel with the biological plantarflexors to stores energy (negative biological power) which is returned later in stance (positive biological power) as done by Collins, Wiggins, and Sawicki [5].

According to our data here, while this approach of storing and returning energy at the ankle can be effective for level ground gaits, at other grades the strategy of immediate storage and return of mechanical energy may not be as effective. Adding a spring in parallel on inclines or declines would likely require an additional biological energy source to inject/extract energy elsewhere in the gait. Another option is to transfer power across joints as depicted in Figure 5E (*i*.*e*., inter-joint transfer). One example is the storage of energy from knee deceleration in late swing and releasing it at the ankle during push-off [2]. From our data, we additionally show that energy storage in the knee during early stance and releasing it at the ankle during push-off becomes increasingly viable with decreasing grade (Figs. 4, 5). A final scenario is that the power from the negative phase could be temporarily stored via battery or clutch and returned at a later time – an approach that has been used within a single gait cycle in foot-ankle prosthesis designs [68, 69]. This last approach, extraction, storage, and then delayed release (Fig. 6F) opens up the possibility to store energy over multiple cycle, perhaps accumulating it, and then return it in a single large burst over a shorter time period to achieve power amplification that may be necessary for on-off accelerations or maximum effort jumps [70].

## Conclusions

Locomotion in the ‘real-world’ involves adjusting speed, changing gait from walk to run and moving up or downhill. The purpose of this study was to characterize changes in lower-limb joint kinetics for walking and running over a range of ground slopes. Specifically, we sought to understand how each joint contributed to total limb positive, negative, and net power output in order to guide development of exoskeleton actuation schemes capable of handling ‘real-world’ mechanical demands. Results of limb-joint level energy analyses motivated us to define three operating modes that exoskeletons could employ: 1) Energy injection: Addition of positive power during positive joint power phases, 2) Energy extraction: Removal of negative power (*i*.*e*., energy harvesting) during negative joint power phase. 3) Energy transfer: extracting energy from one phase and injecting it in another phase at some time later. It’s important to note that we have developed this framework for exoskeletons which operate in parallel with biological muscles and tendons. The guide for development may be different for prostheses which operate in series with biological structures and aim to emulate or fully replace biological joint function [71].

An important next step is to examine whether using biological patterns of joint power output as a ‘road-map’ to apply the three exoskeleton operating modes can improve walking and running performance (*e*.*g*., reduced metabolic cost) on fixed or time varying uphill and downhill slopes.

## Acknowledgements

We would like to thank Moran Gad for his help with the calculation of the inverse dynamics and Karl Zelik for multiple discussions that contributed to aspects of the content in Figure 5.

## Authors’ contributions

GSS, DJF and RR conceived of the study, and designed the experimental protocol. DJF, KZT and RWN carried out experiments. DJF, SM, KZT and RWN analyzed data. RWN drafted the manuscript. GSS, DJF, KZT, RWN, and RR edited the manuscript. All authors gave final approval for publication.

## Availability of data and material

Source data from this study in .mat and .txt format and an associated readme.txt for navigating it are available for download at: http://pwp.gatech.edu/hpl/archival-data-from-publications/

